# Multiple morphological pathways underlie climatic adaptation in North American mustelids

**DOI:** 10.64898/2026.07.24.740590

**Authors:** Chris J. Law

## Abstract

Ecogeographic rules predict that endotherms inhabiting colder environments exhibit larger body sizes and shorter appendages to reduce heat loss. However, these hypotheses have largely been tested using size-based traits, leaving it unclear whether climatic adaptation also alters body proportions that more directly influence surface-area-to-volume ratio. Here, I examined ecogeographic variation in body shape, body size, and relative limb lengths across climatic gradients in three western North American mustelids: American ermine (*Mustela richardsonii*), American mink (*Neogale vison*), and Pacific marten (*Martes caurina*). I then modeled how variation in body shape and size affected surface-area-to-volume ratio. Body size increased in colder climates in all three species, consistently supporting Bergmann’s rule. In contrast, only ermines became stouter in colder climates, whereas body shape remained unchanged in minks and martens. Patterns of appendage variation also differed: ermines and martens exhibited relatively longer limbs in colder climates, contrary to Allen’s rule, whereas only minks exhibited relatively shorter hindlimbs. Modeling showed that increased body size reduced surface-area-to-volume ratio in all species, whereas body shape contributed only in ermines. These findings demonstrate that body shape, body size, and appendage lengths respond independently to climatic gradients, with species-specific ecological and functional constraints shaping thermoregulatory adaptation.

## Introduction

Predicting patterns of phenotypic variation across environmental gradients have long been a central goal for biologists. Ecogeographic patterns such as Bergmann’s rule and Allen’s rule have provided evidence that climate can shape phenotypic evolution [1–3]. Specifically, endotherms in colder climates are predicted to evolve larger body size under Bergmann’s rule and relatively shorter appendages (e.g., limbs, tails, and bills) under Allen’s Rule [4,5]. Both rules stem from the underlying principle that these morphological changes reduce surface-area-to-volume ratio (SA:V) to minimize heat loss in colder climates. Although empirical support for Bergmann’s rule and/or Allen’s rule has been documented across and within diverse birds and mammals [6–11], both rules have traditionally been evaluated using size-based metrics (e.g., body mass, limb length, beak length, tail length). The emphasis on size has led researchers to overlook metrics that capture proportions of the whole body such as body shape, defined here as the ratio between body length and body depth or width.

Changes in overall body shape along an environmental gradient have rarely been tested despite its profound influence on thermoregulation and organismal performance [12–15]. Thermoregulation depends not only on body mass but the distribution of that mass; thus, organisms can have identical body masses but very different body shapes. Because heat exchange is governed by surface area relative to body volume, stout body shapes lose heat more slowly than elongate bodies of similar mass [13,16] and are predicted to occur in colder climates in addition to increased size. However, body shape is also tightly linked to ecological and functional constraints, and selection imposed by locomotion, habitat use, feeding, and phylogenetic history may constrain responses to the climate and environment [17–20]. Consequently, examining variation in body shape along environmental gradients offers an opportunity to evaluate whether thermoregulatory adaptation can overcome functional constraints imposed by ecology as well as investigate its contributions to changes in SA:V across an ecogeographical gradient.

In this study, I investigated ecogeographic variation of body shape, body size, and relative limb lengths in three mustelid species and model how this variation affects changes in SA:V. Mustelids provide an exceptional opportunity to examine this variation because they possess one of the most distinctive body plans among mammals [21–24]. Their elongate bodies facilitate movement through burrows and other confined spaces to capture prey that is typically unavailable to most carnivorans [25,26]. Because this body plan is interpreted as a key innovation underlying the evolutionary and ecological success of the clade [12,27], its functional importance may also limit morphological responses to changing environments. Therefore, if thermoregulatory demands dominate, I predict that individuals that live in colder climates will exhibit larger, stouter body plans with reduced limbs lengths, changes that should also lead to reduction in SA:V. Alternatively, if elongation is maintained by strong functional selection, I predict that climatic adaptation will occur primarily through changes in body size while body shape and relative limb lengths remain similar among the ecogeographical gradient. Thus, body shape changes should have no effect on SA:V. I tested these predictions in three North American mustelids with broad overlapping distributions–the American ermine (*Mustela richardsonii*), the American mink (*Neogale vison*), and the Pacific marten (*Martes caurina*). Together, these species span a wide range of habitats and locomotor ecologies while sharing an elongate body plan: American ermines are highly specialized predators of small mammals that frequently hunt within burrows and beneath snow, conditions that likely favor body elongation; American minks are semi-aquatic generalists that exploit both terrestrial and aquatic prey; and Pacific martens are semi-arboreal predators that use their relative long limbs to hunt in trees and their elongate bodies to also navigate through burrows and snow tunnels [28]. Previous tests of Bergmann’s rule and Allen’s rule have found in mixed patterns for the three species [29–32]; however, these studies examined various populations across all North America and whether environmental factors across the entire continent influenced ecogeographical variation in body size is unknown. Therefore, I limited my sampling to just individuals within the latitudinal range along the west coast of North America from Alaska to southern California.

## Methods

### Morphological data

My dataset consisted of 60 Pacific martens (*Martes caurina*), 45 American ermine (*Mustela richardsonii*), and 36 American minks (*Neogale vison*) from nine natural history museums (see Table S1 for full specimen list). Studied specimens capture the full latitudinal range along the west coast of North America (from Alaska to southern California) of each species. Only specimens with recorded known geographic coordinates were used in this study (Fig. 1). All specimens were fully mature, determined by the closure of exoccipital-basioccipital and basisphenoid-basioccipital sutures on the cranium and full tooth eruption. Because sexual size dimorphism is prevalent in mustelids [33], I only used male specimens to eliminate the effects of sexual dimorphism.

**Fig. 1.**
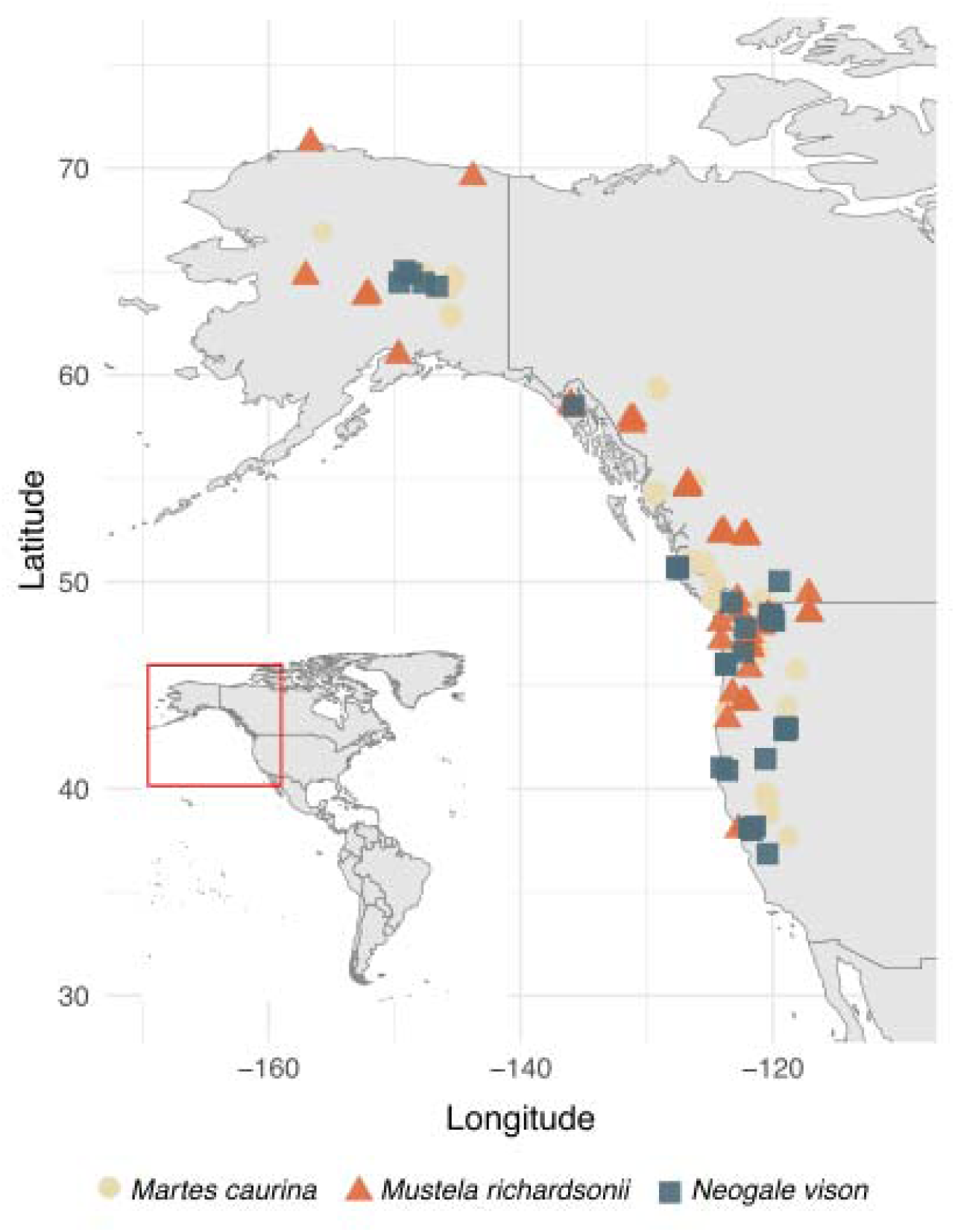
Map of specimens used in this study.

I quantified the body shape of each specimen using the head-body elongation ratio (hbER) [12], which was calculated as the sum of the length of the head and body divided by body depth (head length + body length) / body depth) (Fig. 2). Head length was measured as the condylobasal length of the cranium from the anteriormost point on the premaxilla to the posteriormost point on the surfaces of the occipital condyles; body length was measured by summing the centrum lengths (measured along the ventral surface of the vertebral centrum) of each cervical, thoracic, lumbar, and sacral vertebrae; and body depth was estimated as the average length of the four longest ribs. Each rib was measured as a curve from the end of the capitulum to the point of articulation with the costal cartilage using a flexible measuring tape. High values of hbER suggest a more elongate body shape whereas low values of hbER suggest a stouter body shape. I quantified body size using the geometric mean of the cranial, vertebral, and rib measurements (11th root of the product of our measurements for the cranium, vertebrae, and ribs) [34,35].

**Fig. 2.**
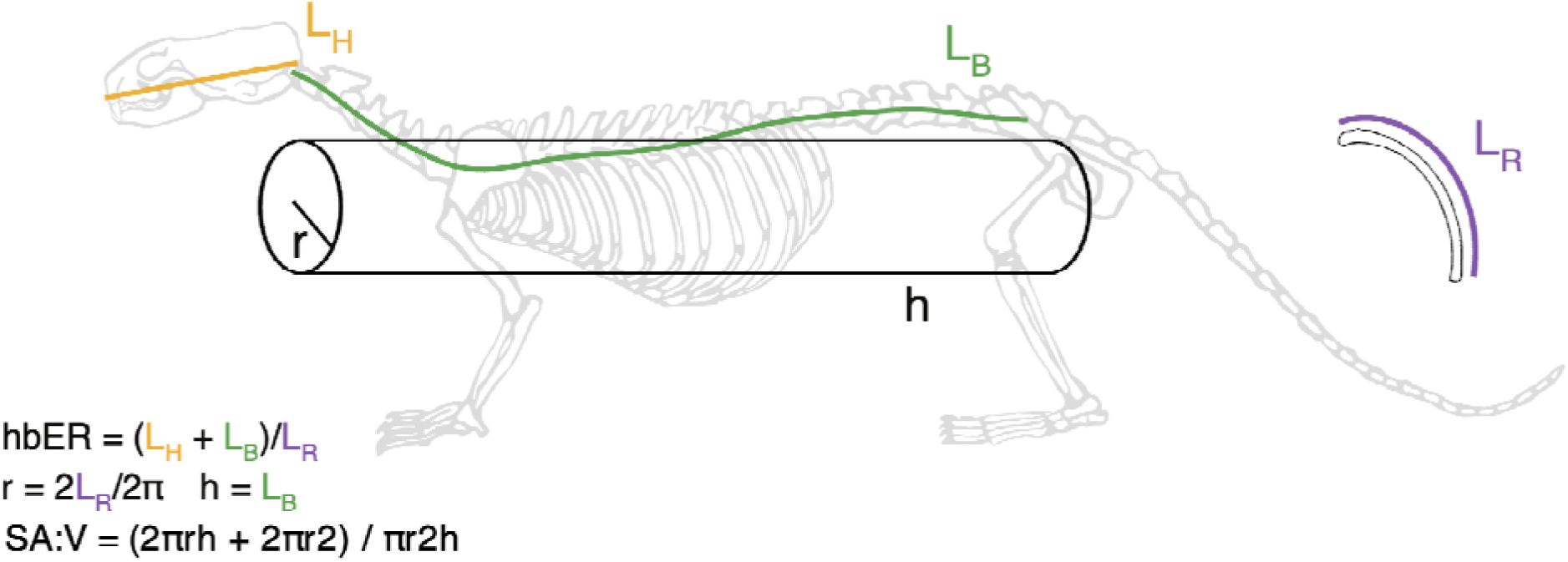
Measurements used to calculate head-body elongation ratio (hbER) and surface-area-to- volume ratio (SA:V) of the body plan. The body shape was modeled as a cylinder, where the radius (r) was estimated based on the curved rib lengths (L_R_) and cylinder height (h) was estimated using body length (L_B_).

Lastly, I measured the lengths of the forelimb and hindlimb. I calculated the forelimb length as the sum of humerus length (measured from the dorsal point of the humeral head to the ventral point of the capitulum) and radius length (measured from the dorsal point of the radial head to the ventral point of the styloid process) and the hindlimb length as the sum of the femur (measured from the dorsal point of the femoral neck to the ventral point of the patellar surface) and tibia lengths (measured from the dorsal point of the intercondylar eminence to the ventral point of the articular surface). I was unable to include the autopodium because most specimens were missing these bones.

### Climatic data

Two proxies of climate were used in this study. Latitudinal data was obtained directly from each specimen and used as the first proxy for climate. Second, I extracted annual mean temperature (BIO1) for each specimen’s geographic coordinates from the WorldClim 2.1 database [36] at a spatial resolution of 10 minutes. Temperature data was extracted using the geodata and terra R packages [37,38]. Latitude and mean annual temperature exhibit a strong relationship (R^2^ = 0.70, P < 0.001), indicating that higher latitudes exhibit lower temperatures.

### Statistical analyses

To investigate Bergmann’s rule, I performed linear regressions to test the relationships between latitude and body size and between latitude and body shape in each species. To investigate Allen’s rule, I performed multiple linear regressions to test the relationships between latitude and forelimb length and between latitude and hindlimb length after accounting for body size in each species. I accounted for body size because there is a strong relationship between body size and limb length (i.e., larger individuals have longer limbs). Partial R^2^ for limb length and body were calculated using the rsq R package [39]. All statistical analyses were performed in R v4.6.0. I then repeated all regression analyses using annual mean temperature.

### Estimation of surface-to-volume ratio

I investigated how changes in body size and body shape influenced surface-area-to-volume ratio (SA:V) by modeling each species as a cylinder (Fig. 2). I used body length as the cylinder height (h) and curved rib length multiplied by two as an estimation of the cylinder’s circumference to estimate the radius (r). I calculated surface-area-to-volume ratio as 2πrh + 2πr^2^ / πr^2^h using empirical body length and curved rib length to model changes in both body size and body shape. To assess the contribution of body size and body shape independently, I calculated SA:V while holding body shape (i.e., r) and body size (i.e., h) constant, respectively. I used mean r and mean h as my constants. I then calculated the percentage change in SA:V between the smallest and largest individuals of each species.

## Results

### Ecogeographic variation in body size, body shape, and relative limb lengths

All relationships using latitude and mean annual temperature exhibited the same pattern. Latitudinal results are reported here and mean annual temperature results are reported in the supplementary materials.

All three mustelids exhibited significant positive relationships between latitude and body size (all P < 0.012; Fig. 3; Table S2). Individuals from higher latitudes exhibited larger body sizes, whereas individuals from lower latitudes exhibited smaller body sizes.

**Fig. 3.**
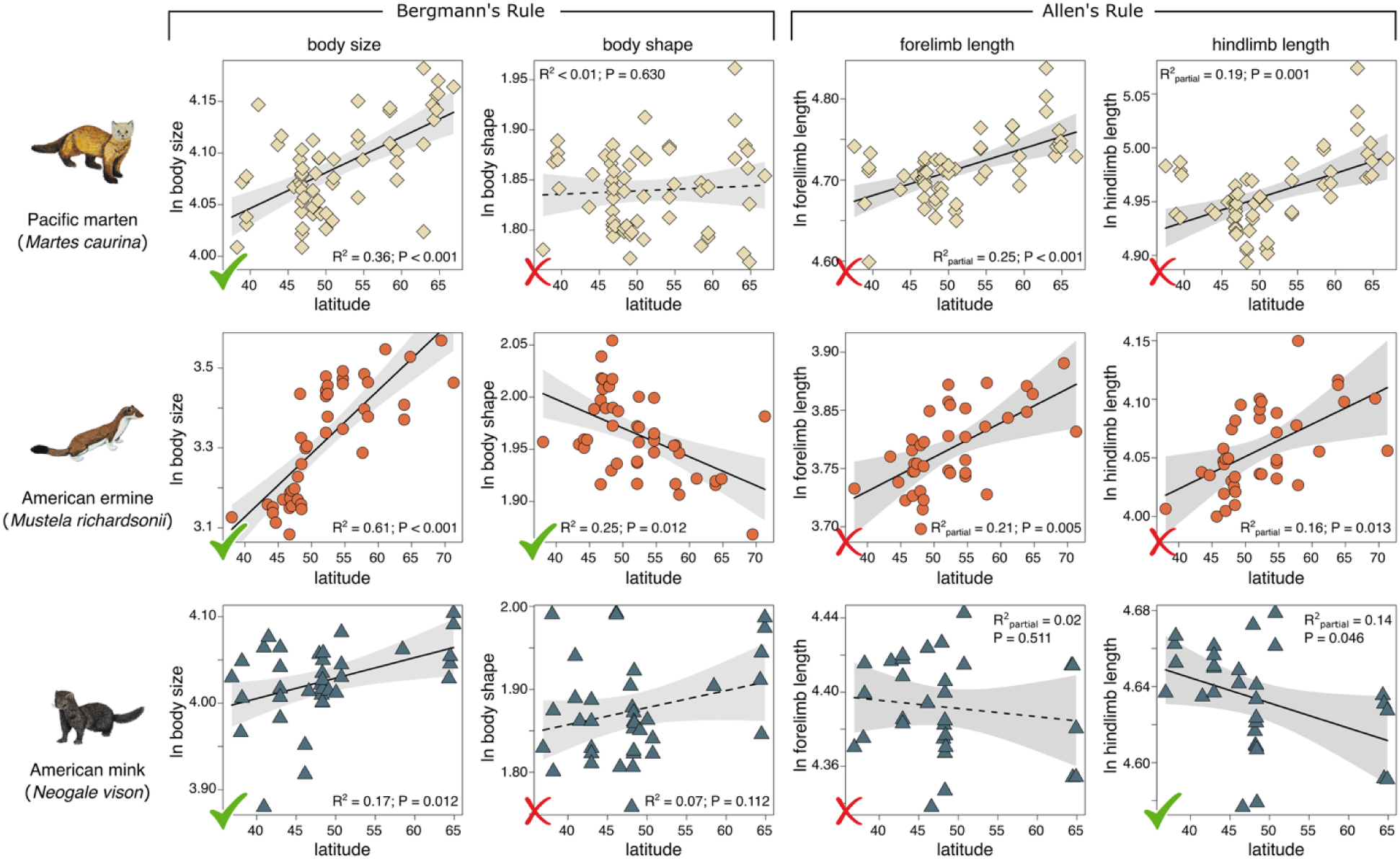
Scatter plots with regressions between latitude and body size, body shape, and limb lengths. Latitude was used as a proxy for climate. The same patterns were found using annual mean temperature (Fig. S1). Solid black lines indicate significant relationships, and dashed black lines indicate non-significant relationships. Gray shaded areas indicate 95% confidence interval of the regression. Green checkmarks indicate support for either Bergmann/Allen’s rule whereas red X’s indicate no support. See Tables S2 and S3 for full ANOVA tables of regression models.

American ermines exhibited a significant negative relationship between latitude and body shape (R^2^ = 0.25, P < 0.001), in which individuals from higher latitudes exhibited stouter body shapes and individuals from lower latitudes exhibited more elongate body shapes (Fig. 3; Table S2). In contrast, there was no significant relationship between latitude and body shape in either Pacific martens (R^2^ = 0.00, P = 0.629) or American minks (R^2^ = 0.07, P = 0.112) (Fig. 3; Table S2).

Both Pacific martens and American ermines exhibited significant positive relationships between latitude and lengths of the forelimb and hindlimb after accounting for body size (all P < 0.013; Fig. 3; Table S3). Individuals from higher latitudes exhibited longer limb lengths in comparison to individuals from lower latitudes. In contrast, American minks did not exhibit a significant relationship between latitude and forelimb length (P = 0.511) but did exhibit a significant negative relationship between latitude and hindlimb length (R^2^ = 0.14, P = 0.046), where individuals from higher latitudes exhibited shorter limb lengths (after accounting for body size) compared to individuals from lower latitudes (Fig. 3; Table S3).

### Estimation of surface-to-volume ratio

Modeling species as cylinders revealed that surface-area-to-volume ratio (SA:V) declined from the smallest to the largest individuals in all three species, although the magnitude of reduction varied substantially. American ermines exhibited the greatest reduction in SA:V at 38.4%. Holding body shape constant indicated that differences in body size alone led to a 26.4% reduction in SA:V, whereas holding body size constant showed that body shape led to a 15.7% reduction, demonstrating that both size and shape influence thermal geometry in this species. American minks exhibited a more modest 16.5% reduction in SA:V, with body shape (9.8%) and body size (7.9%) contributing similarly in SA:V reduction. In contrast, Pacific martens exhibited only a 7.0% reduction in SA:V, which was entirely attributable to increases in body size as holding body shape constant produced the same 7.0% reduction, whereas holding body size constant revealed no differences in SA:V (i.e., 0.0%).

## Discussion

I found that ecogeographic predictions of phenotypic variation are not uniform across the mammalian body plan. Consistent with Bergmann’s rule, body size increased in colder regions in American ermines, American minks, and Pacific martens, suggesting that larger body sizes represent a common response to colder environments across western North American mustelids. In contrast, body shape and appendage proportions exhibited markedly different patterns. Only ermines became stouter in colder regions, whereas body shape remained conserved in mink and martens. Relationships between climate and relative limb lengths were also highly variable: ermines and martens exhibited relatively longer forelimbs and hindlimbs in colder regions–a pattern opposite to predictions of Allen’s rule–whereas only minks exhibited relatively shorter hindlimbs in colder regions. Furthermore, my modeling revealed that all three species reduced surface area-to-volume ratio (SA:V) through different contributions of body size and body shape changes. Together these results demonstrate that maintaining thermoregulatory demands can be achieved through distinct morphological pathways.

The consistent support for Bergmann’s rule across all three species indicates that body size may be the most phenotypically plastic or evolutionarily labile component of the mammalian body plan. Within populations, body size variation reflects changes in environmental factors such as climate and prey availability [40–44]. In mustelids, increased prey availability has led to increased body sizes in populations [29,45,46]. At the evolutionary level, changes in body size are often hypothesized to be the evolutionary path of least resistance [47]. Therefore, body size and size-dependent traits are often the greatest source of phenotypic variation [48–53]. Distinguishing the underlying roles of phenotypic plasticity and evolutionary response of body size geographic variation remains difficult [44]. Nevertheless, the functional goal remains the same regardless of the underlying mechanism: increasing body size reduces SA:V and provides an efficient mechanism for minimizing heat loss. My estimates of SA:V support this interpretation. Across all three species, larger body size was associated with lower SA:V, indicating that increases in body size alone improve thermal efficiency even in the absence of changes in body proportions.

In contrast, patterns between climate and body shape vary among the three species. In minks and martens, climate appears to have less of an impact on body shape variation as predicted. Both species retained similar body shapes across their geographic ranges despite significant increases in body size, whereas only ermines exhibited reduced body elongation at colder regions. The absence of ecogeographical changes in the body shapes of mink and martens demonstrates that larger size alone may be sufficient to achieve thermoregulatory benefits without altering the overall body shape. My SA:V modeling confirms this premise; changes in body shape while holding body size constant show small decreases in SA:V in minks and martens show no change in SA:V in martens. A possible explanation is that functional constraints on body shape may limit responses to climate. Compared to other mammals, minks and martens exhibit elongate body shapes, and like many other mustelids, this body elongation enables them to chase prey in burrows, crevices, and snow tunnels with efficient cost of transport compared to other similar sized mammals [12,25,26]. Selection from locomotion may, therefore, oppose selection from climate and act against shifts towards stouter bodies. Thus, body size appears less constrained than body shape and serves as the evolutionary path of least resistance towards climatic adaptations in these mustelids.

Why ermines alone exhibited reduced body elongation in colder regions remains puzzling. Of the three studied mustelids, ermines are the most specialized in hunting small mammals like voles and mice in burrows, snow tunnels, and other tight crevices [54]. Because their elongate body presumably increases their flexibility and maneuverability to pursue prey in these narrow spaces [12,26], it is counterintuitive to find that ermines are the only mustelid to exhibit stouter bodies. A possible explanation is that as the smallest of the three studied mustelids, ermines may be more susceptible to heat loss in colder climates compared to martens and minks. My modeling revealed that increased body size is not enough to minimize heat loss and proportional changes in the body plan are necessary to reduce SA:V. Specifically, the majority of SA:V reduction in the largest ermines is attributed to body shape changes (26.4% reduction) whereas body size changes contributes only a 15.7% reduction in SA:V. Furthermore, ermines occupy the broadest climatic range of the three species, extending into Arctic and subarctic environments where thermal selection is likely strongest and having bigger, stouter bodies would be beneficial. Although these results suggest that climate may have stronger selective pressure than locomotor demands in ermines, whether reductions in body elongation compromise their ability to hunt within burrows and beneath snow remains to be investigated.

Ecogeographic patterns of limbs further demonstrate that thermoregulatory adaptation cannot be predicted through Allen’s rule. Appendages are typically predicted to become shorter in colder climates to reduce heat loss; however, this expectation was only supported for mink hindlimbs but was contradicted by relatively longer forelimbs and hindlimbs in ermines and martens in colder climates. These results suggest that limb morphology in ermines and martens is shaped by alterative selective pressures beyond thermoregulation. A possible explanation is that relative longer limbs may be influenced by habitat structure rather than climate. Like Pacific martens, American martens (*Martes americana*) also exhibit relatively longer limbs with increasingly colder climates in North America; however, transitions from broadleaf forest to boreal forest was the best explanation for this pattern [32]. Lynch [32] postulated that relatively shorter limbs are more advantageous in exploiting the greater understory complexity that is found broadleaf forests compared to boreal forests. A similar pattern could occur in Pacific martens, in which relatively longer limbs may be more advantageous for vertical climbing in their temperate coniferous forest habitats with limited understory complexity. In ermines, the finding of relatively longer limbs with stouter body shapes is surprising because mustelids typically exhibit shorter limb lengths with increasing body elongation [12]. A potential explanation is that relatively longer limbs may help with locomotion in snow covered landscapes by increasing stride length. Nevertheless, further investigation is clearly needed to uncover potential trade-offs between thermoregulatory and ecological selection that influence relative limb lengths.

Overall, I found that different aspects of the mammalian body plan exhibit varying responses to climatic gradients that may be influenced by species-specific ecological and functional requirements. In general, body size variation but not body shape variation follows predictions of Bergmann’s rule and relative limb lengths exhibited an opposite pattern from predictions of Allen’s rule. Although mammals typically exhibit significant relationships among body size, body shape, and relative limb lengths [12,19,20,23], this study postulates that functional constraints can decouple the evolution (or plasticity) of these traits across ecogeographical gradients such that changes in body size alone can address necessary thermoregulatory demands from colder climates. Thus, species do not need to adhere to both Bergmann’s rule and Allen’s rule to maintain minimum thermoregulator requirements across different climates. Recent work supports this hypothesis, finding that simultaneous adherence to both rules is rare [55,56]. These researchers postulate that the lack of correspondence may be due to species abilities to respond to climate gradient either through changes in body size or appendage size but not both (trade-offs [56]) or through small, coordinated changes in both body size and appendage size instead of a single, larger change in either trait alone (complementarity [55]). Skinner [3] further proposed that assessing all morphological traits together as components of a single thermoregulatory system may be a better approach to assess trade-offs or complementarity in selection from ecogeographical and other ecological and functional factors. My investigation of ecogeographical variation in mustelid body plans follows this framework by using body shape, a prominent feature of vertebrate morphology that may more directly capture an organism’s surface area to volume ratio compared to body size. By explicitly modeling body shape into estimates of SA:V, my analyses provide a more direct assessment of the morphological mechanisms through which climate may influence heat exchange. Unexpectedly however, I found body size and body shape represent independent axes of thermoregulatory adaptation rather than produce a single, “cold climate” body plan. Although all three species converged on increased body size in colder climates, only ermines further reduced SA:V through changes in body shape, whereas mink and martens achieved comparable thermoregulatory outcomes while retaining conserved body proportions. These findings suggest that similar functional objectives can be reached through different morphological pathway and thus highlight the importance of considering whole-body morphology rather than relying solely on body size when evaluating ecogeographic adaptation.

## Ethics

This work did not require ethical approval from a human subject or animal welfare committee.

## Data accessibility

The dataset, R scripts, and Supplementary material will be available online.

## Declaration of AI use

I have not used AI-assisted technologies in creating this article.

## Authors’ contributions

C.J.L.: conceptualization, data curation, formal analysis, funding acquisition, investigation, methodology, project administration, visualization, writing—original draft, writing—review and editing; L.J.H.: funding acquisition, validation, writing—review and editing.

## Conflict of interest declaration

I declare I have no competing interests.

## Funding

Funding was provided by the United States National Science Foundation (awards DBI–2128146 and DEB-2447166) to C.J.L.

## Acknowledgements

I thank the several curators and collection managers for access of specimens held at the Beaty Biodiversity Museum at the University of British Columbia, Burke Museum of Natural History and Culture at the University of Washington, Cal Poly Humboldt Vertebrate Museum, California Academy of Sciences, Field Museum of Natural History, Museum of Natural and Cultural History at the University of Oregon, Museum of Vertebrate Zoology at UC Berkeley, Puget Sound Museum of Natural History, and Royal BC Museum. I also thank Vaibhav Chhaya for the suggestion to model SA:V using a cylinder.

## Notes

### Competing Interest Statement

The authors have declared no competing interest.

