## Supplementary Materials for "Multiple morphological pathways underlie climatic adaptation in North American mustelids"

Chris J. Law

Burke Museum and Department of Biology, University of Washington, Seattle, Washington, USA.

Contains:

Supplementary Results

Fig. S1. Scatter plots with regressions between mean annual temperature and body size, body shape, and limb lengths.

Table S1. List of specimens used in this study.

Table S2. Results of linear regression models to determine how effects of latitude on body size and body shape.

Table S3. Results of multiple regression models to determine how effects of latitude on limb lengths while accounting for body size.

Table S4. Results of linear regression models to determine how effects of annual mean temperature on body size and body shape.

Table S5. Results of multiple regression models to determine how effects of annual mean temperature on limb lengths while accounting for body size.

**Supplementary Results**

All three mustelids exhibited significant positive relationships between annual mean temperature and body size (all P < 0.010; Fig. S1; Table S4). Individuals in lower temperature regions exhibited larger body sizes, whereas individuals from higher temperature regions exhibited smaller body sizes.

American ermines exhibited a significant negative relationship between annual mean temperature and body shape (R^2^ = 0.14, P = 0.011), in which individuals from lower temperature regions exhibited stouter body shapes and individuals from higher temperature regions exhibited more elongate body shapes (Fig. S1; Table S4). In contrast, there was no significant relationship between annual mean temperature and body shape in either Pacific martens (R^2^ = 0.01, P = 0.414) or American minks (R^2^ = 0.05, P = 0.178) (Fig. S1; Table S4).

Both Pacific martens and American ermines exhibited significant positive relationships between annual mean temperature and lengths of the forelimb and hindlimb after accounting for body size (all P < 0.0123; Fig. S1; Table S5). Individuals from lower temperature regions exhibited longer limb lengths in comparison to individuals from higher temperature regions. In contrast, American minks did not exhibit a significant relationship between annual mean temperature and forelimb length (P = 0.455) but did exhibit a significant negative relationship between annual mean temperature and hindlimb length (R^2^_partial_ = 0.18, P = 0.020), where individuals from lower mean temperature regions exhibited shorter limb lengths (after accounting for body size) compared to individuals from higher temperature regions (Fig. S1; Table S5).

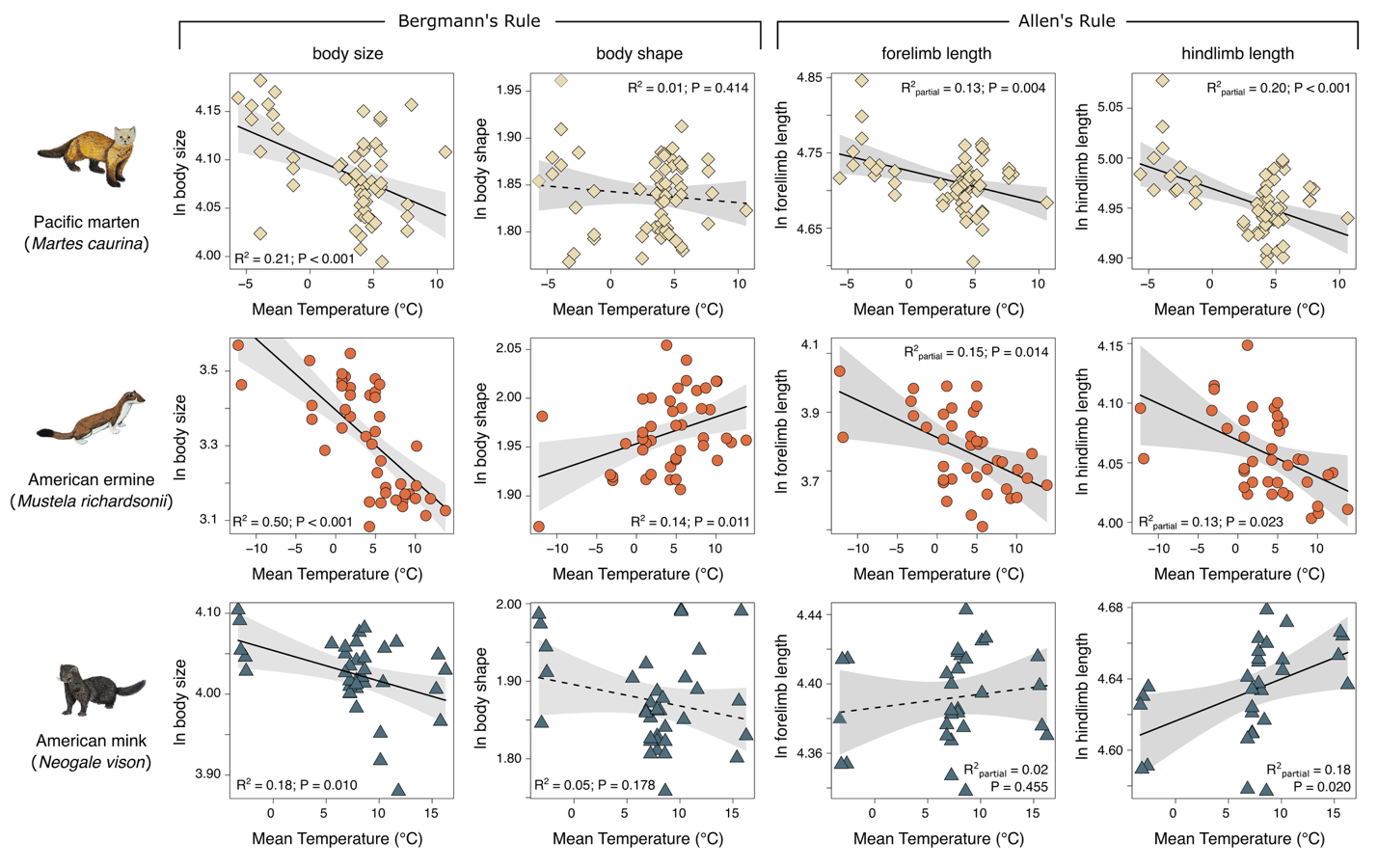

Fig. S1. Scatter plots with regressions between mean annual temperature and body size, body shape, and limb lengths. The same patterns were found using latitude (Fig. 3). Solid black lines indicate significant relationships, and dashed black lines indicate non-significant relationships. Gray shaded areas indicate 95% confidence interval of the regression. See Tables S4 and S5 for full ANOVA tables of regression models.

Table S1. List of specimens used in this study.

| Species | Sex | Catalog Number | State | Latitude | Longitude |
| --- | --- | --- | --- | --- | --- |
| *Martes_caurina* | M | MVZ208655 | CA | 37.56094 | -118.7237 |
| *Martes_caurina* | M | MVZ62774 | CA | 38.928329 | -120.14778 |
| *Martes_caurina* | M | MVZ97308 | CA | 39.4864 | -120.4111 |
| *Martes_caurina* | M | MVZ97307 | CA | 39.4864 | -120.4111 |
| *Martes_caurina* | M | MVZ97309 | CA | 39.4864 | -120.4111 |
| *Martes_caurina* | M | MVZ97311 | CA | 39.6781 | -120.6547 |
| *Martes_caurina* | M | UOMaam13 | OR | 43.6397315 | -123.87132 |
| *Martes_caurina* | M | MVZ83586 | OR | 44.109925 | -118.9412 |
| *Martes_caurina* | M | UWBM12566 | OR | 45.7806 | -118.0917 |
| *Martes_caurina* | M | UWBM52679 | WA | 46.5554 | -121.2614 |
| *Martes_caurina* | M | UWBM52680 | WA | 46.5554 | -121.2614 |
| *Martes_caurina* | M | UWBM36505 | WA | 46.7599 | -121.36 |
| *Martes_caurina* | M | UWBM52674 | WA | 46.7891 | -121.339 |
| *Martes_caurina* | M | UWBM52676 | WA | 46.7891 | -121.339 |
| *Martes_caurina* | M | UWBM34444 | WA | 46.8038 | -121.3179 |
| *Martes_caurina* | M | UWBM34446 | WA | 46.8294 | -121.3684 |
| *Martes_caurina* | M | UWBM52678 | WA | 46.8294 | -121.3684 |
| *Martes_caurina* | M | UWBM36502 | WA | 46.833 | -121.339 |
| *Martes_caurina* | M | UWBM34440 | WA | 46.833 | -121.36 |
| *Martes_caurina* | M | UWBM34441 | WA | 46.833 | -121.2757 |
| *Martes_caurina* | M | UWBM52677 | WA | 46.833 | -121.36 |
| *Martes_caurina* | M | UWBM32593 | WA | 46.915 | -121.6411 |
| *Martes_caurina* | M | UWBM30674 | WA | 46.915 | -121.6411 |
| *Martes_caurina* | M | UWBM52673 | WA | 47.1047 | -121.3538 |
| *Martes_caurina* | M | UWBM33278 | WA | 48.28089 | -120.33261 |
| *Martes_caurina* | M | UWBM33277 | WA | 48.28089 | -120.33261 |
| *Martes_caurina* | M | UWBM33272 | WA | 48.28089 | -120.33261 |
| *Martes_caurina* | M | UWBM33279 | WA | 48.28089 | -120.33261 |
| *Martes_caurina* | M | UWBM33262 | WA | 48.6205 | -119.96496 |
| *Martes_caurina* | M | MVZ12473 | BC | 48.9751 | -124.8 |
| *Martes_caurina* | M | UWBM20609 | BC | 49.0603987 | -120.80465 |
| *Martes_caurina* | M | UBC3497 | BC | 49.064645 | -120.78153 |
| *Martes_caurina* | M | MVZ62824 | BC | 49.3026 | -117.1778 |
| *Martes_caurina* | M | HSU6360 | AK | 58.45671495 | -135.87351 |
| *Martes_caurina* | M | HSU6355 | AK | 58.45671495 | -135.87351 |
| *Martes_caurina* | M | HSU6361 | AK | 58.45671495 | -135.87351 |
| *Martes_caurina* | M | CAS29899 | AK | 62.91388 | -145.53277 |
| *Martes_caurina* | M | CAS29892 | AK | 62.91388 | -145.53277 |
| *Martes_caurina* | M | CAS29894 | AK | 62.91388 | -145.53277 |
| *Martes_caurina* | M | UWBM33290 | AK | 64.1016667 | -145.55917 |
| *Martes_caurina* | M | CAS28697 | AK | 64.2811198 | -146.71209 |
| *Martes_caurina* | M | FMNH151031 | AK | 64.6296692 | -145.25735 |
| *Martes_caurina* | M | FMNH151033 | AK | 64.63006 | -145.25497 |
| *Martes_caurina* | M | UWBM34161 | AK | 64.8333333 | -157.18333 |
| *Martes_caurina* | M | PSM28226 | AK | 64.8377778 | -147.71639 |
| *Martes_caurina* | M | PSM24138 | AK | 66.87322 | -155.67994 |
| *Martes_caurina* | M | RBCM017248 | BC | 54.245126 | -129.24017 |
| *Martes_caurina* | M | RBCM017227 | BC | 54.245126 | -129.24017 |
| *Martes_caurina* | M | RBCM017224 | BC | 54.245126 | -129.24017 |
| *Martes_caurina* | M | RBCM017244 | BC | 54.245126 | -129.24017 |
| *Martes_caurina* | M | RBCM016776 | BC | 50.87 | -125.52 |
| *Martes_caurina* | M | RBCM016777 | BC | 50.87 | -125.52 |
| *Martes_caurina* | M | RBCM017031 | BC | 51.029954 | -126.59014 |
| *Martes_caurina* | M | RBCM017024 | BC | 51.029954 | -126.59014 |
| *Martes_caurina* | M | RBCM017020 | BC | 50.010612 | -124.57079 |
| *Martes_caurina* | M | RBCM017011 | BC | 50.010612 | -124.57079 |
| *Martes_caurina* | M | RBCM016549 | BC | 59.38327 | -129.11132 |
| *Martes_caurina* | M | RBCM016550 | BC | 59.38327 | -129.11132 |
| *Martes_caurina* | M | RBCM016528 | BC | 59.38327 | -129.11132 |
| *Martes_caurina* | M | RBCM016541 | BC | 54.882333 | -126.0921 |
| *Mustela_richardsonii* | M | MVZ191001 | CA | 38.0353414 | -122.79672 |
| *Mustela_richardsonii* | M | UWBM72863 | OR | 43.4259 | -123.4751 |
| *Mustela_richardsonii* | M | HSU4345 | OR | 44.18699286 | -122.20083 |
| *Mustela_richardsonii* | M | HSU4349 | OR | 44.18699286 | -122.20083 |
| *Mustela_richardsonii* | M | PSM10319 | OR | 44.6333 | -123.2928 |
| *Mustela_richardsonii* | M | MVZ190246 | WA | 45.70303972 | -121.8833 |
| *Mustela_richardsonii* | M | UWBM56648 | WA | 46.6861 | -121.81326 |
| *Mustela_richardsonii* | M | UWBM56152 | WA | 46.6897192 | -121.70856 |
| *Mustela_richardsonii* | M | UWBM33273 | WA | 46.7900192 | -121.8917 |
| *Mustela_richardsonii* | M | UWBM57518 | WA | 46.7900192 | -121.8917 |
| *Mustela_richardsonii* | M | UWBM30243 | WA | 47.0175 | -121.88861 |
| *Mustela_richardsonii* | M | UWBM39367 | WA | 47.0328 | -124.1628 |
| *Mustela_richardsonii* | M | UWBM32601 | WA | 47.4256 | -121.7756 |
| *Mustela_richardsonii* | M | UWBM72864 | WA | 47.9179 | -124.28903 |
| *Mustela_richardsonii* | M | UWBM32594 | WA | 47.94554 | -120.76758 |
| *Mustela_richardsonii* | M | UWBM33273 | WA | 48.28089 | -120.33261 |
| *Mustela_richardsonii* | M | UWBM80453 | WA | 48.4152 | -120.3796 |
| *Mustela_richardsonii* | M | UWBM41278 | WA | 48.4208333 | -122.65 |
| *Mustela_richardsonii* | M | UWBM76594 | WA | 48.4845 | -117.0853 |
| *Mustela_richardsonii* | M | UWBM76605 | WA | 48.4868 | -117.101 |
| *Mustela_richardsonii* | M | UBC19518 | BC | 49.0928537 | -122.75801 |
| *Mustela_richardsonii* | M | MVZ62800 | BC | 49.3026 | -117.1778 |
| *Mustela_richardsonii* | M | UWBM41237 | BC | 52.1417 | -122.1417 |
| *Mustela_richardsonii* | M | UWBM41238 | BC | 52.1417 | -122.1417 |
| *Mustela_richardsonii* | M | UWBM41239 | BC | 52.1417 | -122.1417 |
| *Mustela_richardsonii* | M | UWBM41240 | BC | 52.1417 | -122.1417 |
| *Mustela_richardsonii* | M | UBC254 | BC | 52.4 | -124.03333 |
| *Mustela_richardsonii* | M | UBC256 | BC | 52.4 | -124.03333 |
| *Mustela_richardsonii* | M | UBC255 | BC | 52.4 | -124.03333 |
| *Mustela_richardsonii* | M | MVZ31025 | BC | 57.9 | -131.1667 |
| *Mustela_richardsonii* | M | MVZ31026 | BC | 57.9 | -131.1667 |
| *Mustela_richardsonii* | M | HSU5139 | AK | 58.45671495 | -135.87351 |
| *Mustela_richardsonii* | M | HSU5140 | AK | 58.45671495 | -135.87351 |
| *Mustela_richardsonii* | M | PSM16974 | AK | 61.069 | -149.7653 |
| *Mustela_richardsonii* | M | PSM18459 | AK | 63.89 | -152.23583 |
| *Mustela_richardsonii* | M | PSM18460 | AK | 63.89 | -152.23583 |
| *Mustela_richardsonii* | M | UWBM33489 | AK | 64.8 | -157.2 |
| *Mustela_richardsonii* | M | MVZ123992 | AK | 69.45972222 | -143.75194 |
| *Mustela_richardsonii* | M | MVZ134451 | AK | 71.29055556 | -156.78861 |
| *Mustela_richardsonii* | M | RBCM014873 | BC | 54.686948 | -126.69592 |
| *Mustela_richardsonii* | M | RBCM014879 | BC | 54.686948 | -126.69592 |
| *Mustela_richardsonii* | M | RBCM014877 | BC | 54.686948 | -126.69592 |
| *Mustela_richardsonii* | M | RBCM014885 | BC | 54.686948 | -126.69592 |
| *Mustela_richardsonii* | M | RBCM014891 | BC | 54.686948 | -126.69592 |
| *Mustela_richardsonii* | M |  |  | 57.6507286 | -131.1667 |
| *Neogale_vision* | M | MVZ32188 | CA | 36.8589 | -120.455 |
| *Neogale_vision* | M | MVZ129726 | CA | 37.9942818 | -121.57817 |
| *Neogale_vision* | M | MVZ126839 | CA | 38.142389 | -121.48783 |
| *Neogale_vision* | M | MVZ44140 | CA | 38.1514 | -121.9717 |
| *Neogale_vision* | M | HSU8550 | CA | 40.91519557 | -123.62101 |
| *Neogale_vision* | M | HSU1262 | CA | 40.92025428 | -124.08596 |
| *Neogale_vision* | M | MVZ80764 | CA | 41.5106493 | -120.49671 |
| *Neogale_vision* | M | UWBM41791 | OR | 42.9718 | -118.9144 |
| *Neogale_vision* | M | UWBM41783 | OR | 42.9718 | -118.9144 |
| *Neogale_vision* | M | UWBM41784 | OR | 42.9718 | -118.9144 |
| *Neogale_vision* | M | UWBM41803 | OR | 42.9718 | -118.9144 |
| *Neogale_vision* | M | UWBM41789 | OR | 42.9718 | -118.9144 |
| *Neogale_vision* | M | MVZ89538 | OR | 46.1157 | -123.83 |
| *Neogale_vision* | M | MVZ89537 | OR | 46.1157 | -123.83 |
| *Neogale_vision* | M | UWBM32604 | WA | 46.6020317 | -122.2739 |
| *Neogale_vision* | M | UWBM32612 | WA | 47.908983 | -122.10883 |
| *Neogale_vision* | M | UWBM32108 | WA | 48.1379407 | -119.9022 |
| *Neogale_vision* | M | UWBM32117 | WA | 48.2193541 | -120.115 |
| *Neogale_vision* | M | UWBM32114 | WA | 48.3061918 | -120.115 |
| *Neogale_vision* | M | UWBM32116 | WA | 48.3061918 | -120.115 |
| *Neogale_vision* | M | UWBM32121 | WA | 48.3061918 | -120.115 |
| *Neogale_vision* | M | UWBM32110 | WA | 48.320182 | -120.1211 |
| *Neogale_vision* | M | UWBM32113 | WA | 48.3346547 | -120.1211 |
| *Neogale_vision* | M | UWBM32115 | WA | 48.3780727 | -120.1211 |
| *Neogale_vision* | M | UWBM32112 | WA | 48.407018 | -120.1211 |
| *Neogale_vision* | M | UBC019497 | BC | 49.09858314 | -123.1778 |
| *Neogale_vision* | M | UBC019499 | BC | 50.05072735 | -119.39789 |
| *Neogale_vision* | M | UBC260 | BC | 50.720821 | -127.49666 |
| *Neogale_vision* | M | UBC258 | BC | 50.720821 | -127.49666 |
| *Neogale_vision* | M | UBC223 | BC | 50.720821 | -127.49666 |
| *Neogale_vision* | M | HSU6386 | AK | 58.45671495 | -135.87351 |
| *Neogale_vision* | M | PSM28215 | AK | 64.3164782 | -146.66237 |
| *Neogale_vision* | M | PSM24955 | AK | 64.4766 | -147.708 |
| *Neogale_vision* | M | CAS29891 | AK | 64.4783 | -149.5768 |
| *Neogale_vision* | M | PSM28211 | AK | 64.891111 | -148.81028 |
| *Neogale_vision* | M | PSM28210 | AK | 64.8983 | -149.1973 |

Table S2. Results of linear regression models to determine how effects of latitude on body size and body shape.

| Pacific marten (*Martes caurina*) | | | | |  |  |  |
| --- | --- | --- | --- | --- | --- | --- | --- |
|  | *body size ~ latitude* | | SS | DF | R^2^ | F | P |
|  |  | latitude | 0.04 | 1 | 0.36 | 32.91 | <0.001 |
|  |  | Residuals | 0.07 | 58 |  |  |  |
|  | *body shape ~ latitude* | | SS | DF | R^2^ | F | P |
|  |  | latitude | 0.00 | 1 | <0.01 | 0.23 | 0.630 |
|  |  | Residuals | 0.09 | 58 |  |  |  |
| American ermine (*Mustela richardsonii*) | | | | | |  |  |
|  | *body size ~ latitude* | | SS | DF | R^2^ | F | P |
|  |  | latitude | 0.55 | 1 | 0.61 | 66.89 | <0.001 |
|  |  | Residuals | 0.36 | 43 |  |  |  |
|  | *body shape ~ latitude* | | SS | DF | R^2^ | F | P |
|  |  | latitude | 0.02 | 1 | 0.25 | 14.40 | <0.001 |
|  |  | Residuals | 0.05 | 43 |  |  |  |
| American mink (*Neogale vison*) | | | |  |  |  |  |
|  | *body size ~ latitude* | | SS | DF | R^2^ | F | P |
|  |  | latitude | 0.01 | 1 | 0.17 | 7.04 | 0.012 |
|  |  | Residuals | 0.06 | 34 |  |  |  |
|  | *body shape ~ latitude* | | SS | DF | R^2^ | F | P |
|  |  | latitude | 0.01 | 1 | 0.07 | 2.67 | 0.112 |
|  |  | Residuals | 0.12 | 34 |  |  |  |

Table S3. Results of multiple regression models to determine how effects of latitude on limb lengths while accounting for body size.

| Pacific marten (*Martes caurina*) | | |  |  |  |  |  |
| --- | --- | --- | --- | --- | --- | --- | --- |
|  | *forelimb length ~ latitude + body size* | | SS | DF | R^2^_partial_ | F | P |
|  |  | latitude | 0.02 | 1 | 0.25 | 18.87 | <0.001 |
|  |  | body size | 0.04 | 1 | 0.41 | 38.43 | <0.001 |
|  |  | Residuals | 0.06 | 57 |  |  |  |
|  | *hindlimb length ~ latitude + body size* | | SS | DF | R^2^_partial_ | F | P |
|  |  | latitude | 0.01 | 1 | 0.19 | 13.52 | 0.001 |
|  |  | body size | 0.04 | 1 | 0.45 | 46.50 | <0.001 |
|  |  | Residuals | 0.05 | 56 |  |  |  |
| American ermine (*Mustela richardsonii*) | | |  |  |  |  |  |
|  | *forelimb length ~ latitude + body size* | | SS | DF | R^2^_partial_ | F | P |
|  |  | latitude | 0.01 | 1 | 0.21 | 8.88 | 0.005 |
|  |  | body size | 0.28 | 1 | 0.91 | 351.57 | <0.001 |
|  |  | Residuals | 0.03 | 36 |  |  |  |
|  | *hindlimb length ~ latitude + body size* | | SS | DF | R^2^_partial_ | F | P |
|  |  | latitude | 0.01 | 1 | 0.16 | 6.91 | 0.013 |
|  |  | body size | 0.36 | 1 | 0.92 | 399.00 | <0.001 |
|  |  | Residuals | 0.03 | 36 |  |  |  |
| American mink (*Neogale vison*) | | |  |  |  |  |  |
|  | *forelimb length ~ latitude + body size* | | SS | DF | R^2^_partial_ | F | P |
|  |  | latitude | 0.00 | 1 | 0.02 | 0.44 | 0.511 |
|  |  | body size | 0.03 | 1 | 0.68 | 47.54 | <0.001 |
|  |  | Residuals | 0.02 | 27 |  |  |  |
|  | *hindlimb length ~ latitude + body size* | | SS | DF | R^2^_partial_ | F | P |
|  |  | latitude | 0.00 | 1 | 0.14 | 4.40 | 0.046 |
|  |  | body size | 0.03 | 1 | 0.66 | 53.45 | <0.001 |
|  |  | Residuals | 0.02 | 27 |  |  |  |

Table S4. Results of linear regression models to determine how effects of annual mean temperature on body size and body shape.

| Pacific marten (*Martes caurina*) | | |  |  |  |  |  |
| --- | --- | --- | --- | --- | --- | --- | --- |
|  | *body size ~ temperature* | | SS | DF | R^2^ | F | P |
|  |  | temperature | 0.025 | 1 | 0.21 | 15.828 | 0.000 |
|  |  | Residuals | 0.091 | 58 |  |  |  |
|  | *body shape ~ temperature* | | SS | DF | R^2^ | F | P |
|  |  | temperature | 0.001 | 1 | 0.01 | 0.678 | 0.414 |
|  |  | Residuals | 0.092 | 58 |  |  |  |
| American ermine (*Mustela richardsonii*) | | | |  |  |  |  |
|  | *body size ~ temperature* | | SS | DF | R^2^ | F | P |
|  |  | temperature | 0.452 | 1 | 0.50 | 42.566 | 0.000 |
|  |  | Residuals | 0.457 | 43 |  |  |  |
|  | *body shape ~ temperature* | | SS | DF | R^2^ | F | P |
|  |  | temperature | 0.010 | 1 | 0.14 | 7.140 | 0.011 |
|  |  | Residuals | 0.057 | 43 |  |  |  |
| American mink (*Neogale vison*) | | |  |  |  |  |  |
|  | *body size ~ temperature* | | SS | DF | R^2^ | F | P |
|  |  | temperature | 0.013 | 1 | 0.18 | 7.352 | 0.010 |
|  |  | Residuals | 0.059 | 34 |  |  |  |
|  | *body shape ~ temperature* | | SS | DF | R^2^ | F | P |
|  |  | temperature | 0.007 | 1 | 0.05 | 1.893 | 0.178 |
|  |  | Residuals | 0.122 | 34 |  |  |  |

Table S5. Results of multiple regression models to determine how effects of annual mean temperature on limb lengths while accounting for body size.

| Pacific marten (*Martes caurina*) | | |  |  |  |  |  |
| --- | --- | --- | --- | --- | --- | --- | --- |
|  | *forelimb length ~ temperature + body size* | | SS | DF | R^2^_partial_ | F | P |
|  |  | temperature | 0.010 | 1 | 0.13 | 8.822 | 0.004 |
|  |  | ln body size | 0.069 | 1 | 0.51 | 59.806 | <0.001 |
|  |  | Residuals | 0.066 | 57 |  |  |  |
|  | *hindlimb length ~ temperature + body size* | | SS | DF | R^2^_partial_ | F | P |
|  |  | temperature | 0.012 | 1 | 0.20 | 14.078 | <0.001 |
|  |  | ln body size | 0.058 | 1 | 0.55 | 69.329 | <0.001 |
|  |  | Residuals | 0.047 | 56 |  |  |  |
| American ermine (*Mustela richardsonii*) | | |  |  |  |  |  |
|  | *forelimb length ~ temperature + body size* | | SS | DF | R^2^_partial_ | F | P |
|  |  | temperature | 0.006 | 1 | 0.15 | 6.657 | 0.014 |
|  |  | ln body size | 0.341 | 1 | 0.92 | 407.751 | <0.001 |
|  |  | Residuals | 0.030 | 36 |  |  |  |
|  | *hindlimb length ~ temperature + body size* | | SS | DF | R^2^_partial_ | F | P |
|  |  | temperature | 0.005 | 1 | 0.13 | 5.609 | 0.023 |
|  |  | ln body size | 0.435 | 1 | 0.93 | 465.652 | <0.001 |
|  |  | Residuals | 0.034 | 36 |  |  |  |
| American mink (*Neogale vison*) | | |  |  |  |  |  |
|  | *forelimb length ~ temperature + body size* | | SS | DF | R^2^_partial_ | F | P |
|  |  | temperature | 0.000 | 1 | 0.02 | 0.575 | 0.455 |
|  |  | ln body size | 0.034 | 1 | 0.64 | 47.215 | <0.001 |
|  |  | Residuals | 0.020 | 27 |  |  |  |
|  | *hindlimb length ~ temperature + body size* | | SS | DF | R^2^_partial_ | F | P |
|  |  | temperature | 0.004 | 1 | 0.18 | 6.086 | 0.020 |
|  |  | ln body size | 0.036 | 1 | 0.68 | 57.875 | <0.001 |
|  |  | Residuals | 0.017 | 27 |  |  |  |
